# Amphibians in the Brazilian Cerrado: diversity, research effort and conservation

**DOI:** 10.1101/2020.02.13.945618

**Authors:** Enne Gomes Alves, Fernando Mayer Pelicice

## Abstract

In order to investigate the relationship among amphibian diversity, research effort and conservation status across the Brazilian Cerrado, we carried out a scientometric analysis. Scientific publications were searched in international databases, totaling 72 studies, with 177 taxa recorded. Most studies occurred in the Southeast and Midwest regions of the country, followed by Northeast and North. The first two regions summed almost all taxa; controlling for sampling effort, however, northern regions appeared more diverse. Moreover, northern regions still preserve much of the original vegetation, while savanna in Southeast and Midwest regions has been significantly suppressed. These differences indicate particular geographical needs: immediate conservation efforts to protect diversity and restore savannas in Southeast and Midwest regions; and basic research effort (inventory) to uncover biodiversity in northern Cerrado areas.

## Introduction

The Cerrado biome is the largest savanna ecosystem in South America, and originally covered about 2 million km^2^; c.a. 20% of Brazilian territory (Ratter *et al.* 1997; Klink & Machado 2005; Sano *et al.* 2007). Over the past decades, however, the biome has been significantly reduced by human activities (Verdade *et al.* 2010), especially livestock raising, agriculture and hydroelectric plants. According to Sano *et al*. (2008), 80 million hectares are used by humans, and agriculture and pasture cover 37% of the biome. Yet, Brazilian Cerrado still preserves a rich biodiversity (Klink & Machado 2005), making it a hotspot for conservation (Myers *et al*. 2000).

Cerrado’s biodiversity is expressed in different taxonomic groups, including amphibians (Amphibia). From 877 amphibians species recorded in Brazil (SBH 2010), 134 are found in the Cerrado savanna, being 42 endemic (Colli *et al*. 2002; Klink & Machado 2005; Bastos 2007). Amphibians, particularly anurans, have been studied for decades in the Cerrado biome (Sazima & Bokermann 1978; Duellman 1979), but basic questions remain, such as geographical distribution, natural history, ecology, and total diversity (Diniz-Filho *et al*. 2004, Silvano & Segalla 2005; Leite *et al*. 2008). The need for knowledge is critical, considering that the biome is changing rapidly and amphibians have particular biological features. They have a permeable skin, and their life cycle depends on aquatic and terrestrial habitats, so the group is especially vulnerable to minimum environmental disturbances. Currently, amphibians are globally threatened with extinction (Verdade *et al.* 2010).

Because the biome is geographically extensive, heterogeneous and human influence is unevenly distributed across regions (Sano *et al.* 2008), it is essential to examine existing knowledge about the issue; i.e. the spatial distribution of research effort and known amphibian diversity. This understanding may guide future research and, coupled with information on conservation status of savannas, may assist management and conservation decisions. The present study, therefore, examined scientific publications that investigated amphibians in the Brazilian Cerrado in order to analyze the relationship between research effort, diversity and conservation status of the biome. Using scientometric and meta-analysis approaches, we investigated (i) total amphibian species richness, (ii) the geographical distribution of diversity and (iii) research effort, and (iv) the relationship between these variables and the conservation status of the biome (% remaining area).

## Material and Methods

### Data sampling

A scientometric analysis (August 31, 2011) was performed to collect scientific publications that investigated amphibians in the Cerrado biome. We consulted datasets provided by Web of Science (www.isiknowledge.com), Scopus (www.scopus.com) and Scielo (www.scielo.org), complemented with articles published by Check List journal (www.checklist.org.br). The first two index global scientific literature, while Scielo is concerned mainly with Latin American. Check List was searched because it is specialized in publishing local species inventories. All articles were collected using the following combination of words: “anura*cerrado”, “anura*savanna*Brazil”, “amphibia*cerrado” and “amphibia*savanna*Brazil”.

The search resulted in 101 articles. Eleven were not analyzed because they were not available (n = 2) or did not focus on amphibians (n = 9). The remainder (n =90) was then divided among literature review (n = 8), meta-analysis (n = 10) and research (n = 72). The first two categories were not considered and only research papers (species description, biology and inventory) were analyzed here.

From each article, we recorded species composition and the region where the study took place: South (S), Southeast (SE), Midwest (MW), Northeast (NE) or North (N). Considering each study as an independent observational unit, we built a matrix containing all recorded species (columns: presence/absence) and individual studies (rows). Species nomenclature followed AMNH (2011).

### Data analysis

We calculated the number of studies (research effort) and taxa recorded (known diversity) for each region. To compare species richness among regions, we used taxa rarefaction curves, controlling for research effort. Curves were generated by randomizing (500 times) the species x effort matrix, using the software Estimates version 8.2.0 (Colwell, 2009). Non-parametric estimators were used to estimate total taxa richness expected in the biome. Following Brose (2002), we chose Chao 2, Jacknife 1 and 2.

To investigate the relationship between research effort, species richness and conservation status of the Cerrado biome, we calculated the original biome area (ha) and the current remaining cover (ha and %) – data from Sano et al. (2007). With these data, we investigated qualitatively the relationship between remaining area (%) versus species richness and research effort. In this case, richness was controlled for the minimum research effort among regions (n = 3), using rarefaction results.

## Results

Of the 72 studies retained, 51 focused on populations (e.g. taxonomic description, biology), while 21 carried out assemblage inventories. Research inventories were less common in all regions, ranging from 24% in Midwest to 40% in Northeast regions (Figure 1A). In total, most studies (89%) took place in Southeast and Midwest, followed by Northeast and North regions (Figure 1A). No study was recorded in South (two articles did not report location).

**Figure 1.**
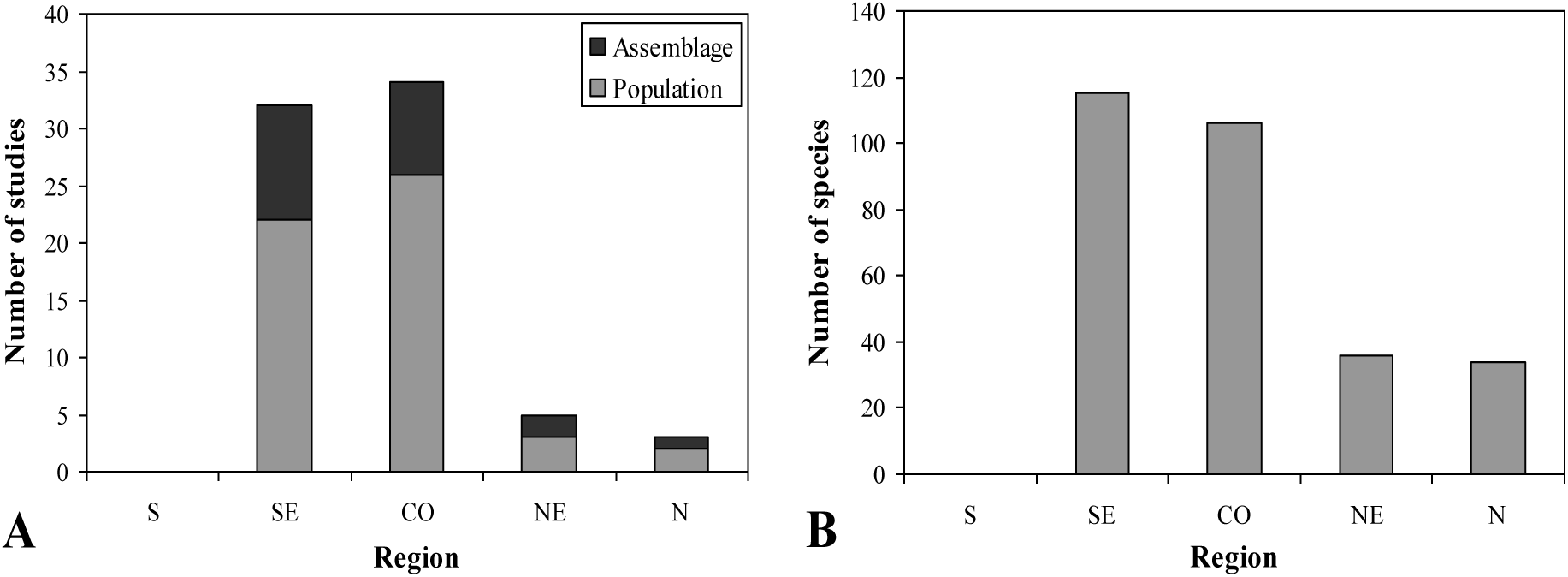
Number of studies that investigated amphibians in Brazilian Cerrado (population studies or assemblage inventories; A), and number of species recorded in these studies (B). Regions: South (S), Southeast (SE), Midwest (MW), Northeast (NE) and North (N).

We recorded 177 taxa, distributed in 15 families (Table S1). The most representative were Hylidae (S = 76) and Leptodactylidae (S =27). The Southeast (S = 115) and Midwest (S = 106) regions summed most taxa (Figure 1B), followed by North (S = 36) and Northeast (S = 34). The rarefaction curve showed no tendency to stabilization (Figure 2A), indicating that new species will be added with additional studies. In fact, total taxa richness estimated by non-parametric estimators was much higher than the observed value (Chao 2 = 270; Jacknife 1 = 261; Jacknife 2 = 308). Lack of stabilization also characterized each region (Figure 2B). In addition, when sampling effort is controlled, the North region appeared as the most diverse, indicating richer local assemblages.

**Figure 2.**
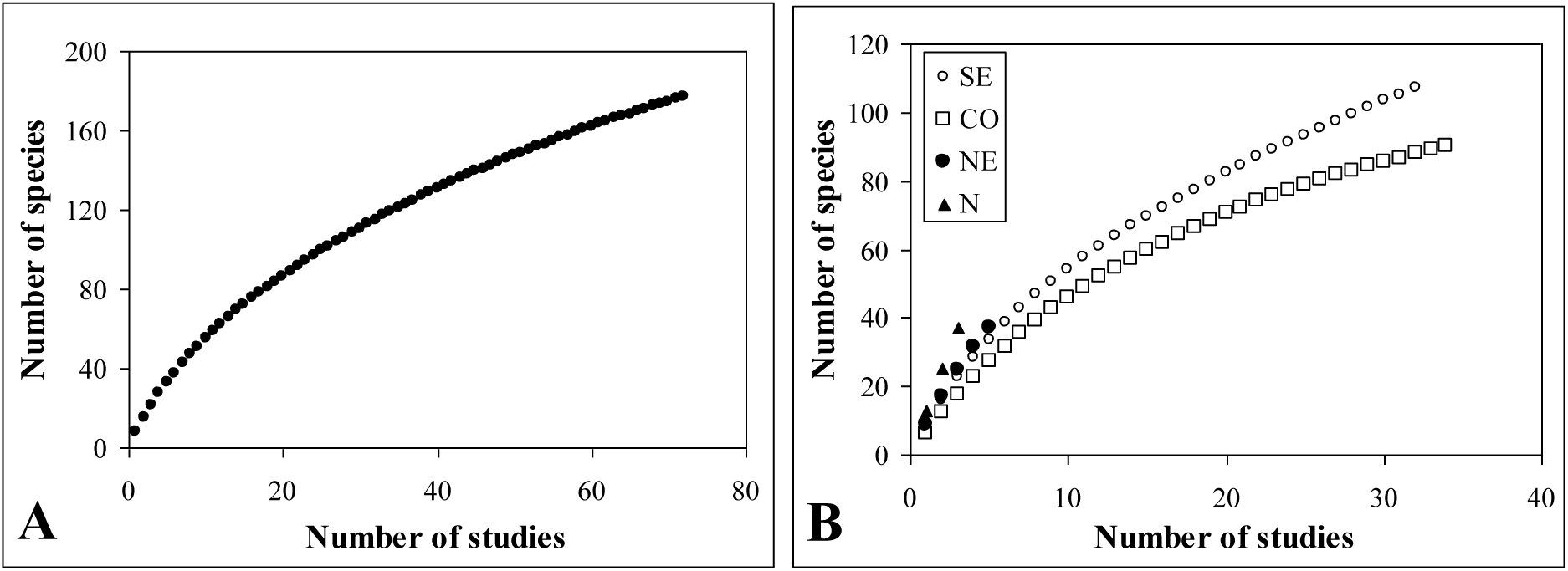
Total amphibian species richness (rarefaction curves) recorded in the Brazilian Cerrado (A) and in each region of the biome (B). See Figure 1 for region’s codes.

Originally, Midwest had the largest Cerrado area, followed by Northeast, Southeast and North regions (Table 1). The smallest area occurred in the South region (Table 1). At present, large proportions of remnant vegetation are found in northern regions (> 80%), while Midwest, Southeast and South lost at least half of their original cover (Table 1). Regions with high percentage of remaining area had higher taxa richness (Figure 3A). On the other hand, regions with small percentage of remaining area received more research effort (Figure 3B) – except for the South region, where no study was registered.

**Table 1.** Original area covered by the Cerrado biome (ha) and current remaining area (ha and %) in different Brazilian regions. Data based on Sano et al. (2007).

| Region | Original area<br>(ha) | Remaining area<br>(ha) | Remaining area<br>(%) |
| --- | --- | --- | --- |
| South | 374,257 | 118,692 | 31.7 |
| Southeast | 41,226,482 | 18,873,589 | 45.8 |
| Midwest | 91,009,437 | 45,595,960 | 50.1 |
| Northeast | 45,593,730 | 38,554,184 | 84.6 |
| North | 25,090,246 | 20,251,786 | 80.7 |

**Figure 3.**
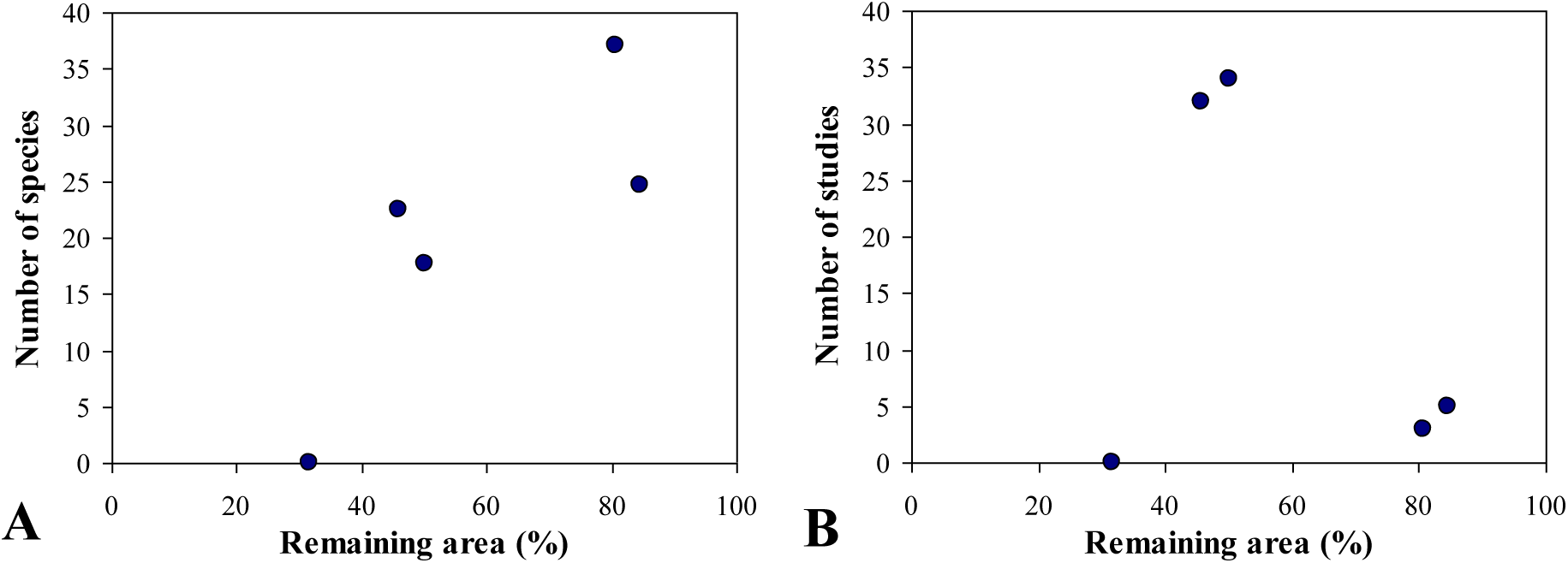
Relationship between remaining savanna area (%) versus number of species recorded (A) and research effort (B). See Figure 1 for region’s codes.

## Discussion

Scientometric analysis retrieved 72 studies that investigated amphibians in the Brazilian Cerrado, basically anurans. Taken together, these studies reported 177 taxa, a high value if compared to the literature (see discussion below). We also noted geographical differences, since most recorded taxa and research effort occurred in Southeast and Midwest regions of Brazil. It should be noted, however, that regions with high proportions of original savanna (e.g. North and Northeast), although less studied, showed high biodiversity potential.

Total taxa richness (177) was higher than reported by previous studies, such as Colli *et al*. (2002) (110 species), Bastos (2007) (142 species) and Klink & Machado (2005) (150 species). The existence of synonyms or incorrect identification could explain this increase, but taxa identification and organization followed AMNH (2011). Actually, this result is probably associated with increased research effort in the biome, considering that 82% of analyzed articles were published after 2005. We emphasize, therefore, that amphibian diversity is underestimated in the biome. For example, scientometry is a technique that sample indexed literature, so there must be other studies that were not covered here. Moreover, as suggested by rarefaction curves and richness estimators, new species will be recorded with increasing research effort, especially in Southeast, Northeast and North regions. Finally, we point out that some taxa were identified at the genus level, so they may cluster different species.

Research effort and current known diversity concentrated in Southeast and Midwest Brazil. These regions, where major research institutes and universities are located, hold nearly all knowledge about amphibians in the Cerrado biome. In fact, the existence of meta-analysis studies (e.g. Diniz-Filho *et al*. 2005, 2006 and 2007) strongly indicates that there is enough data for synthesis and large-scale evaluations. In addition, these studies have also recorded higher species richness in Southeast and Midwest regions, and here we show a possible connection with research effort. In contrast, we observed that northern regions received lower research effort and, consequently, had lower inventoried diversity. Only three studies, for example, were reported in the North (Tocantins state). Diversity in these regions is underestimated and rarefaction curves strongly suggested this trend: controlling for sampling effort, the North region showed the highest taxa richness, indicating rich local diversity. Moreover, northern regions still preserve much of the original savanna (Sano *et al*. 2008), so they must hold endemic and unknown species. Therefore, to better understand amphibian diversity patterns in Brazilian Cerrado (e.g. total diversity and spatial distribution), research effort, especially inventories, must increase in northern regions of the country.

An important finding was the mismatch between the amount of knowledge and conservation status. In the case, regions with considerable proportion of the original biome (North and Northeast) had much lower research effort and, consequently, lower diversity known. Differently, regions that received more research effort (Southeast and Midwest), with relatively well-known anurans, have lost considerable Cerrado areas. This trend emphasizes the existence of different geographical demands: the need for effective conservation measures in southern and Midwest regions, in order to preserve/restore degraded areas and prevent further habitat loss; and basic research effort (inventory) in northern regions. Finally, it is important to highlight that no study was recorded in the South region. In fact, these Cerrado remnants have received little research attention (Uhlmann et al., 1998). Comparatively, South Brazil had smaller original savanna area, but habitat loss has been intense. This situation (weak research effort, coupled with intense human influence), stress the urgent need for fauna inventories and conservation measures.

Brazilian Cerrado is a hotspot for biodiversity conservation (Myers *et al*. 2000). The biome, however, has been progressively reduced (Klink & Machado 2005), especially due to agriculture and the construction of large dams. This trend, coupled with the fact that amphibian diversity is declining at global scale, stress the need for conservation measures that protect amphibians in this biome. Conservation success will depend, in turn, on high quality information, enabling the identification of pressing environmental issues, diversity patterns and eventual conflicts in land/water usage. The present study, by examining the scientific literature, indicated priority areas of research and conservation, and showed that current demands are not distributed evenly across the biome: there are geographical mismatches among research effort, existing knowledge, and conservation status. These results, together with other studies (Diniz-Filho *et al*. 2005, 2006 and 2007), may guide future decisions concerning the allocation of effort to management, conservation and research activities.

## Acknowledgement

We thank Adriana B. Andrade (UFT) for suggestions, and the Núcleo de Estudos Ambientais (Neamb) and Universidade Federal do Tocantins (UTF) for providing infrastructure and logistic.

