## Supplementary material for "Amphibians in the Brazilian Cerrado: diversity, research effort and conservation": Table S1

### Supplementary file

Table S1. Amphibians recorded in the Brazilian Cerrado, according to scientometric results. Regions: South (S), Southeast (SE), Midwest (MW), Northeast (NE) and North (N).

| | S | SE | MW | N | NE |
|---|---|---|---|---|---|
| **ANURA** | | | | | |
| **Aromobatidae** | | | | | |
| *Allobates goyanus* | | | x | | |
| **Brachycephalidae** | | | | | |
| *Eleutherodactylus* sp. (gr. *lacteus*) | | | | | x |
| *Ischnocnema izecksohni* | | x | | | |
| *Ischnocnema juipoca* | | x | | | |
| *Ischnocnema penaxavantinho* | | x | | x | |
| **Bufonidae** | | | | | |
| *Bufo* sp. 1 (gr. *granulosus*) | | | x | | |
| *Bufo* sp. 2 (gr. *granulosus*) | | | x | | |
| *Melanophryniscus fulvoguttatus* | | | x | | |
| *Rhinella bergi* | | | x | | |
| *Rhinella cerradensis* | | x | | | |
| *Rhinella granulosa* | | x | x | | |
| *Rhinella major* | | | x | | |
| *Rhinella margaritifera* | | x | | x | |
| *Rhinella mirandaribeiroi* | | | x | | |
| *Rhinella ocellata* | | | x | x | x |
| *Rhinella ornata* | | x | x | x | |
| *Rhinella pombali* | | x | | | |
| *Rhinella rubescens* | | x | x | x | |
| *Rhinella schneideri* | | x | x | | |
| *Rhinella veredas* | | x | x | | x |

| | | | | |
|---|---|---|---|---|
| **Centrolenidae** | | | | |
| *Hyalinobatrachium* sp. | x | | | |
| *Hyalinobatrachium* sp. (aff. *eurygnatum*) | x | | | |
| *Vitreorana uranoscopa* | x | | x | x |
| **Ceratophryidae** | | | | |
| *Ceratophrys cranwelli* | | x | | |
| *Ceratophrys joazeirensis* | | x | | |
| **Craugastoridae** | | | | |
| *Haddadus binotatus* | x | | | |
| **Cycloramphidae** | | | | |
| *Odontophrynus americanus* | | x | | |
| *Odontophrynus cultripes* | | x | | |
| *Odontophrynus* sp. | | x | | |
| *Proceratophrys boiei* | | | x | |
| *Proceratophrys cristiceps* | | x | | |
| *Proceratophrys goyana* | x | x | | x |
| *Proceratophrys* sp. | x | | | |
| *Proceratophrys strussmannae* | | x | | |
| *Proceratophrys vielliardi* | | x | | |
| **Dendrobatidae** | | | | |
| *Adelphobates galactonotus* | | | x | |
| *Ameerega berohoka* | | x | | |
| *Ameerega braccata* | | x | | |
| *Ameerega flavopicta* | x | x | | |
| **Hylidae** | | | | |
| *Aplastodiscus arildae* | x | | | |
| *Bokermannohyla alvarengai* | x | | | |
| *Bokermannohyla circumdata* | x | | | |
| *Bokermannohyla martinsi* | x | x | x | x |
| *Bokermannohyla nanuzae* | x | | | |

| | | | | |
|---|---|---|---|---|
| *Bokermannohyla pseudopseudis* | | | x | |
| *Bokermannohyla saxicola* | x | | | |
| *Dendropsophus cruzi* | x | x | | |
| *Dendropsophus elegans* | x | | | |
| *Dendropsophus elianeae* | x | | x | |
| *Dendropsophus jimi* | | x | | |
| *Dendropsophus leucophyllatus* | | x | x | x |
| *Dendropsophus melanargyreus* | x | x | | x |
| *Dendropsophus microcephalus* | | x | | |
| *Dendropsophus minutus* | x | x | | |
| *Dendropsophus nanus* | x | x | x | x |
| *Dendropsophus rubicundulus* | | x | | |
| *Dendropsophus seniculus* | x | x | x | x |
| *Dendropsophus soaresi* | x | | | |
| *Hypsiboas albomarginatus* | x | | | |
| *Hypsiboas albopunctatus* | x | x | | |
| *Hypsiboas caingua* | x | | | |
| *Hypsiboas cipoensis* | x | | | x |
| *Hypsiboas crepitans* | | | x | x |
| *Hypsiboas faber* | x | | x | x |
| *Hypsiboas geographicus* | x | | | |
| *Hypsiboas goianus* | x | | | |
| *Hypsiboas lundii* | x | x | | |
| *Hypsiboas multifaciatus* | x | | x | x |
| *Hypsiboas polytaenius* | x | | | |
| *Hypsiboas pulchellus* | x | | | |
| *Hypsiboas punctatus* | x | x | | x |
| *Hypsiboas raniceps* | x | x | | |
| *Hypsiboas wavrini* | x | | | |
| *Itapotihyla langsdorffii* | x | x | x | x |
| *Phasmahyla jandaia* | | x | | |
| *Phyllomedusa ayeaye* | | x | | |
| *Phyllomedusa azurea* | | x | | |
| *Phyllomedusa boliviana* | | x | | |
| *Phyllomedusa burmeisteri* | x | | | |

| | | | | |
|---|---|---|---|---|
| *Phyllomedusa camba* | | x | | |
| *Phyllomedusa hypochondrialis* | x | x | | x |
| *Phyllomedusa sauvagii* | | x | | x |
| *Phyllomedusa tetraploidea* | x | x | | |
| *Pseudis limellum* | | x | | |
| *Pseudis paradoxa* | | x | | |
| *Pseudis platensis* | x | x | | |
| *Pseudis tocantins* | x | | | |
| *Scinax acuminatus* | | x | | |
| *Scinax berthae* | | x | | |
| *Scinax canastrensis* | | x | | |
| *Scinax cardosoi* | x | | | |
| *Scinax catharinae* | x | | x | |
| *Scinax centralis* | | x | | |
| *Scinax curicica* | | x | | x |
| *Scinax duartei* | x | | | |
| *Scinax eurydice* | | x | x | |
| *Scinax fuscomarginatus* | x | x | | |
| *Scinax fuscovarius* | x | x | | x |
| *Scinax hiemalis* | x | | | |
| *Scinax luizotavioi* | | x | | |
| *Scinax machadoi* | x | x | x | x |
| *Scinax nasicus* | x | x | | |
| *Scinax nebulosus* | | | x | |
| *Scinax perereca* | x | | | |
| *Scinax rizibilis* | x | | | |
| *Scinax ruber* | x | x | | |
| *Scinax similis* | | x | | |
| *Scinax* sp. | | | | x |
| *Scinax* sp.1 (aff. *perereca*) | x | | | |
| *Scinax* sp.2 (gr. *catharinae*) | | | | x |
| *Scinax* sp.3 | | | x | |
| *Scinax squalirostris* | x | | | |
| *Scinax x-signatus* | x | | | |
| *Trachycephalus mambaiensis* | x | | | |

| | | | | |
|---|---|---|---|---|
| *Trachycephalus venulosus* | x | x | x | x |
| **Hylodidae** | | | | |
| *Crossodactylus bokermanni* | x | | | |
| *Crossodactylus caramaschii* | x | | | |
| *Crossodactylus trachystomus* | | | | x |
| *Hylodes uai* | x | | | |
| **Leiuperidae** | | | | |
| *Eupemphix nattereri* | x | x | | |
| *Physalaemus albifrons* | x | | | |
| *Physalaemus albonotatus* | x | x | | |
| *Physalaemus biligonigerus* | x | x | | |
| *Physalaemus centralis* | x | x | | |
| *Physalaemus cuvieri* | x | x | | |
| *Physalaemus evangelistai* | x | x | x | x |
| *Physalaemus marmoratus* | x | | x | |
| *Physalaemus olfersii* | x | | | |
| *Physalaemus* sp. | x | | | |
| *Pleurodema diplolister* | x | | | |
| *Pleurodema fuscomaculata* | x | | | |
| *Pseudopaludicola canga* | x | | | x |
| *Pseudopaludicola falcipes* | x | x | | |
| *Pseudopaludicola mystacalis* | x | x | | x |
| *Pseudopaludicola saltica* | | x | | |
| *Pseudopaludicola* sp. | x | | | |
| *Pseudopaludicola ternetzi* | | | x | |
| **Leptodactylidae** | | | | |
| *Leptodactylus andreae* | | x | | |
| *Leptodactylus bokermanni* | x | | | x |
| *Leptodactylus bufonius* | | x | | |
| *Leptodactylus chaquensis* | x | x | | |
| *Leptodactylus cunicularius* | | x | | |
| *Leptodactylus diptyx* | | x | | |

| | | | | |
|---|---|---|---|---|
| *Leptodactylus elenae* | x | x | | |
| *Leptodactylus furnarius* | x | x | | |
| *Leptodactylus fuscus* | x | x | x | |
| *Leptodactylus hylaedactylus* | x | x | | |
| *Leptodactylus jolyi* | | x | | |
| *Leptodactylus labyrinthicus* | x | x | | |
| *Leptodactylus latrans* | x | x | x | x |
| *Leptodactylus lauramiriamae* | x | | | |
| *Leptodactylus macrosternum* | x | x | x | x |
| *Leptodactylus marmoratus* | x | x | x | |
| *Leptodactylus martinezi* | | x | | |
| *Leptodactylus mystaceus* | x | x | x | |
| *Leptodactylus mystacinus* | x | x | | |
| *Leptodactylus petersi* | | x | | |
| *Leptodactylus podicipinus* | x | x | x | x |
| *Leptodactylus sertanejo* | x | x | | |
| *Leptodactylus* sp. | x | x | x | |
| *Leptodactylus* sp. (= *Adenomera*) | x | | | |
| *Leptodactylus syphax* | x | x | | |
| *Leptodactylus troglodytes* | x | | | |
| *Leptodactylus vastus* | | x | | |
| **Microhylidae** | | | | |
| *Chiasmocleis albopunctata* | x | x | | |
| *Chiasmocleis mehelvi* | x | | | |
| *Dermatonotus muelleri* | x | x | | |
| *Elachistocleis bicolor* | | x | | x |
| *Elachistocleis ovalis* | x | x | x | x |
| *Elachistocleis piauiensis* | x | x | x | |
| *Elachistocleis* sp.1 | | x | | |
| *Elachistocleis* sp.2 | | x | | |
| **Strabomantidae** | | | | |
| *Barycholos ternetzi* | x | x | | |
| *Oreobates heterodactylus* | x | | | |

| | | |
|---|---|---|
| *Pristimantis fenestratus* | x | |
| *Pristimantis* sp. | x | |
| **GYMNOPHIONA** | | |
| **Caeciliidae** | | |
| *Siphonops paulensis* | | x |
